# Revisiting Reddy: A DLBCL Do-over

**DOI:** 10.1101/2023.11.21.567983

**Authors:** Kostiantyn Dreval, Manuela Cruz, Christopher Rushton, Nina Liuta, Houman Layegh Mirhosseini, Callum Brown, Ryan D. Morin, the GAMBL consortium

## Abstract

The 2017 study by Reddy *et al* described the comprehensive characterization of somatic drivers of diffuse large B-cell lymphoma using whole exome sequencing.^1^ After additional large studies relying on exome or whole genome sequencing were published, several oddities unique to the Reddy results have emerged. Seeking to explain the discrepancies, we reanalyzed their data using established open-source pipelines. This revealed thousands of mutations that could not be independently reproduced by these pipelines and a larger set of high-quality mutations that were not reported by Reddy. This caused an artificial under-representation of the mutation prevalence in many genes including clinically relevant hot spots affecting *EZH2* and *CD79B*. More generally, the study had an under-representation of mutations in DLBCL genes that disproportionately affected genes known to have the highest mutation rates. The missing variants and the spurious variants can be attributed to distinct problems with the analytical approaches employed in that study. Our analysis also identified strong associations between mutations and patient outcome including *TP53, KMT2D* and *PIM1*, which were not found in the Reddy study. Overall, we demonstrate that this combination of errors influenced many of the central novel findings from their study rendering their results largely non-replicable. The full results of our analyses are included as supplemental items as a resource for other researchers with an interest in the genetics of B-cell lymphomas.

## Introduction

Reddy *et al* described their use of whole exome sequencing to study the interplay between somatic alterations and clinical outcomes in Diffuse Large B-cell lymphoma (DLBCL).^1^ Boasting 1001 patient samples, this still represents the largest sequencing effort applied to any B-cell lymphoma to date. The authors reported the comprehensive identification of 150 genetic drivers (referred to here as Reddy gene list) including 27 genes not previously attributed to this neoplasm. Assuming the data quality was sufficient and that appropriate analytical methods were applied, the study had the potential to exhaustively identify all commonly mutated genes in DLBCL. This analysis could also inform on the existence and frequency of mutation hot spots – which can represent gain-of-function mutations – in the DLBCL patient population. The authors used their mutation and gene expression information to train a “Genomic Risk Model”, which they claimed could be useful for risk stratification.

Since the arrival of the Reddy study, multiple large exome and whole genome sequencing-based analyses of DLBCL independently identified the genetic drivers of DLBCL and a set of genetic subgroups defined by combinations of these.^2,3^ Several of the genes that are consistently mutated in only one of these subgroups, such as *NOTCH1*, *GRHPR*, *ITPKB*, *ZFP36L1* and *DTX1*, were conspicuously absent from the Reddy gene list, rendering them fully redacted from their results. Oddly, another set of genes that had been previously reported to represent DLBCL drivers were also absent from the Reddy list, including *HIST1H1C* and *ID3*. With an ultimate goal to resolve discrepancies in the literature and clarify the mutation patterns in lymphoid cancers, our team is leading a meta-analysis of all exome- and genome-based studies from B-cell neoplasms. The genomics of mature B-cell lymphomas (GAMBL) project uses a combination of established open-source Snakemake^4^ pipelines and custom analyses in our R package (GAMBLR).

Using the GAMBL results, we attempted to reproduce the findings from Reddy *et al*. This uncovered a multitude of discrepancies that can only be explained by several serious problems in the analytical approaches used to perform the original analysis. Thousands of variants reported by Reddy were not detected by our variant calling approach and these tended to have marginal support in the primary data. One of the most egregious issues we identified was a systematic error that caused the loss of hundreds of hot spot mutations including those in clinically relevant genes such as *EZH2*^5^ and *CD79B*^6^ leading them to report unrealistically low mutation prevalence for these genes. We hope that these analyses and results presented herein will provide a valuable resource for other researchers attempting to use the Reddy data.

## Results

### Reddy or not?

We identified candidate somatic simple somatic mutations in each of the 999 exomes that were available in the European Genome-Phenome Archive (**Methods**). These variants were then restricted to genes that have been ascribed to DLBCL biology either in the Reddy study or more recent studies (237 genes; **Supplemental Table S1**). Indels were not considered in subsequent analyses, leaving 13451 SNVs from GAMBL. These were divided into three sets according to whether they were also reported in the original study, yielding a set of GAMBL-only (6675), Intersect (3568), and Reddy-only (3145) SNVs (**Supplemental Table S2**). Within the 150 genes reported by Reddy, there are 3817 GAMBL-only mutations. The genes with the highest number of Reddy-only variants include *PIM1* (118), *KMT2D* (115), *BCL2* (71), *ARID1A* (71), and *ARID1B* (68). Interestingly, some of these same genes harbored the most GAMBL-only variants such as *KMT2D* (214), *PIM1* (452) and *BCL2* (270). This is consistent with the notion that genes with more *bona fide* mutations somehow had a higher likelihood of being erroneously assigned mutations in the Reddy analysis. Although this led to elevated mutation incidence in these genes, the mutations were distinct from those identified by GAMBL and, importantly, not necessarily in the same patients.

We also noted that the Reddy analysis completely ignored the relevance of splice site mutations, which explains some of the discrepancies. However, only a small minority of the remaining GAMBL-only variants represent splice site mutations, leaving the bulk of the discrepancies unresolved by this difference. When considering individual patients, the number of discrepancies between the results varied considerably, with one patient having as many as 36 discordant mutations (28 GAMBL-only and 8 Reddy-only)(**Table 1**). Strikingly, of the 999 patients that were re-analyzed, 183 (18%) had *zero* concordant mutations and only 8 patients showed 100% concordance between the two analyses (**Supplemental Table S3**). There were 385 patients with more Reddy-only variants than GAMBL-only variants and 475 patients showing the converse pattern. Alignments showing the reads surrounding each mutation can be accessed on the companion website (https://www.bcgsc.ca/downloads/morinlab/GAMBL/Reddy/igv_reports/).

**Table 1.** Examples of patients with large discrepancies between Reddy and GAMBL. Each GAMBL ID links to an IGV report showing the primary data supporting each variant. *If the embedded hyperlinks in the table are not functional, each example can be found on this page:* https://www.bcgsc.ca/downloads/morinlab/GAMBL/Reddy/igv_reports/

| Sample ID | GAMBL ID | GAMBL-only variants | Intersect | Reddy-only variants |
| --- | --- | --- | --- | --- |
| 2766 | Reddy_2766T | 1 | 0 | 13 |
| 3429 | Reddy_3429T | 5 | 0 | 3 |
| 3585 | Reddy_3585T | 0 | 0 | 9 |
| 830 | Reddy_830T | 15 | 0 | 6 |
| 3834 | Reddy_3834T | 28 | 17 | 7 |
| 2153 | Reddy_2153T | 23 | 9 | 5 |
| 2213 | Reddy_2213T | 17 | 6 | 0 |
| 2566 | Reddy_2566T | 7 | 0 | 0 |

In theory, our analytical approach, which involves four variant callers, is more conservative than the one used in the original study. It was surprising that the number of variants in the GAMBL results was higher than those unique to Reddy. Nonetheless, some of the Reddy-only variants may not have been identified here due to technical nuances of each variant caller. The validity of mutations unique to either analysis cannot be experimentally determined without access to the samples. In lieu of this, we can only evaluate the quality of the variant calls objectively using the primary data. Using a simplified approach to that established previously,^7^ we assigned a group of curators to review the exome data in integrative genomics viewer IGV^8^ for all the three sets, with curators blinded to the origin of each variant to avoid unconscious bias. In each review, variants were assigned a rating on a 5-point scale with 1 reserved for variants having the minimal support (one molecule) and 5 representing variants with the best support (**Figure 1A**). Each variant was reviewed by at least one curator and the average rating for each variant was calculated. Comparing the distributions of these ratings across the three sets revealed a clear enrichment for variants with dubious support (ratings of 1 or 2) in the Reddy-only list relative to those unique to GAMBL. Supporting the quality of the variants from GAMBL, the Intersect list had a nearly identical score distribution to the GAMBL-only set (**Figure 1B**).

**Figure 1.**
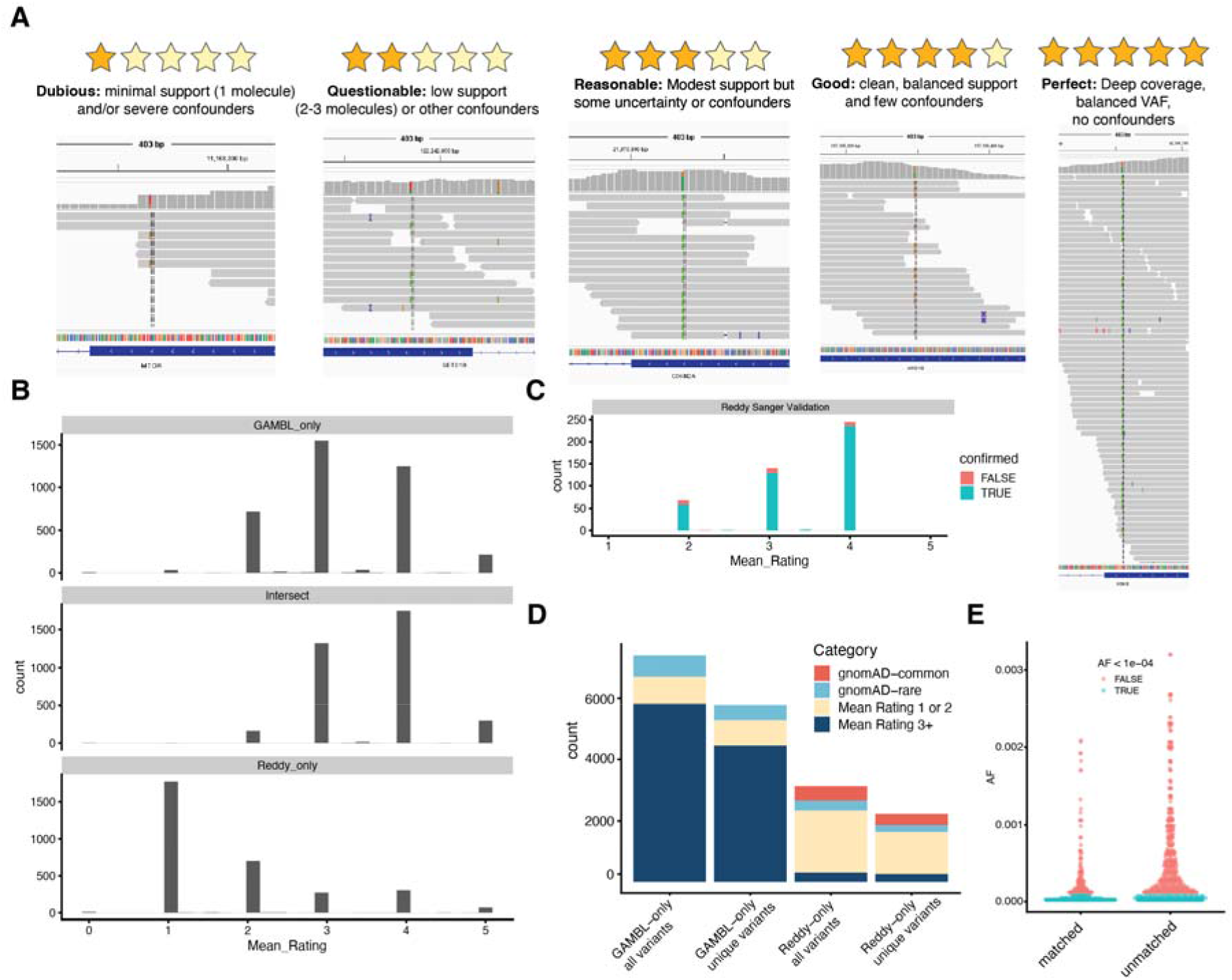
Objective evaluation of variant quality with visual inspection. (A) The 5-point scale we used to score each variant is explained along with representative examples from each of the scoring categories. All scoring was performed using a custom web interface implemented using the Shiny R package. All curators were blinded to the origin of each variant when they performed this scoring. (B) For each variant we calculated the mean value across the curators. The three histograms show the distribution of these values within the three sets of variants. The GAMBL-only set (top) are SNVs detected in Reddy genes by the GAMBL pipeline but not reported by Reddy et al. The Intersect set (middle) are those variants reported by Reddy and identified by our pipeline. The Reddy-only set (bottom) are those variants that were unique to the Reddy analysis. (C) Although the majority of Reddy-only variants were rated as 1 or 2, those selected for Sanger validation in the Reddy study were predominantly rated as 3 or 4. (D) Variants in each set were annotated using gnomAD AF and categorized as gnomAD-common (AF>0.0001); gnomAD-rare (present in gnomAD but with a lower AF); within the lowest two ratings in our scale (1-2) or in the upper 3. This bar chart separately shows the number of variants with and without collapsing them to the non-redundant alleles. (E) The variants from patients that had matched normal exomes were separated from the remaining unmatched patients. The number of variants in gnomAD and their AF are plotted with the common variants shown in salmon (AF > 0.0001).

The Reddy study reported a 90% concordance based on Sanger sequencing, which implies a relatively high accuracy of their variant calls. The observation that almost 50% of their variant calls were not reproduced by our analysis implies the actual accuracy was, in fact, much lower. Interestingly, we noticed that the variants selected for validation by Sanger sequencing in the Reddy study tended to be assigned high ratings based on our blinded review (**Figure 1C**). Based on the distribution of ratings in their full set of variants, we conclude that those variants selected for Sanger sequencing were not representative of the overall quality of their results. Whether by design or a side-effect of their selection of variants favoring certain genes, their validation experiment was heavily enriched for variants with strong support in the primary data. The rate of true positives in the Reddy results can be estimated instead based on discordance with GAMBL, which places it closer to 53% than the 90% reported in that study.

A subset of the Reddy-only variants were assigned higher ratings, which implies they are well-supported in the data but, for some reason, were not identified by our pipeline. One possible explanation is the persistence of common germline variants in the Reddy results. Aiming to enrich for somatic mutations as much as possible in the absence of a matched normal, the GAMBL pipeline automatically removes variants matching common gnomAD alleles, which we consider as having overall allele fraction (AF) > 0.0001. Indeed, 478 of the Reddy-only set appear to be missing from the GAMBL results because they had gnomAD AF values that exceed our threshold (**Figure 1D**). After excluding the gnomAD variants from the Reddy-only list, nearly all the remaining variants have ratings of 1 or 2, implying they had very low support in the primary data. The existence of common, and likely germline, variants in the Reddy-only list was surprising because matched normal exomes were supposedly sequenced from 400 of the 1001 patients (though the data was not released with the tumor exomes). After identifying the tumors with matched normal and separating the 999 analyzed tumors based on matched/unmatched status, we saw that common gnomAD variants remained in both groups. The gnomAD variants were significantly more abundant among the unmatched samples, which is consistent with the authors having removed some (but not all) germline variants from these cases using the matched exome data (**Figure 1E**).

### These aren’t the genes you’re looking for

The Reddy study presented a core set of 150 DLBCL genes that they identified using a combination of previously published data and their exomes. The discrepancies between the Reddy results and the GAMBL results were unevenly distributed across these genes. The effect of this was an artificial inflation of mutation frequency in some genes and an underestimate of mutation incidence in others (**Figure 2A-B**). Even when the GAMBL results are used, the mutation incidence of DLBCL genes is lower in the Reddy cohort than the collection of DLBCLs from more recent studies with a few notable exceptions (**Figure 2C**). Some of these variants could be attributed to contamination of the exome libraries with PCR product (amplicon). In particular, candidate hot spots in *PIK3CD*, *STAT5B* and *ATM* could be explained by this issue (**Figure 3**). Mutations in *PIK3CD* were first attributed to DLBCL by the same group but there were no mutations at the artefactual hot spot in their earlier description of this pattern.^9^

**Figure 2.**
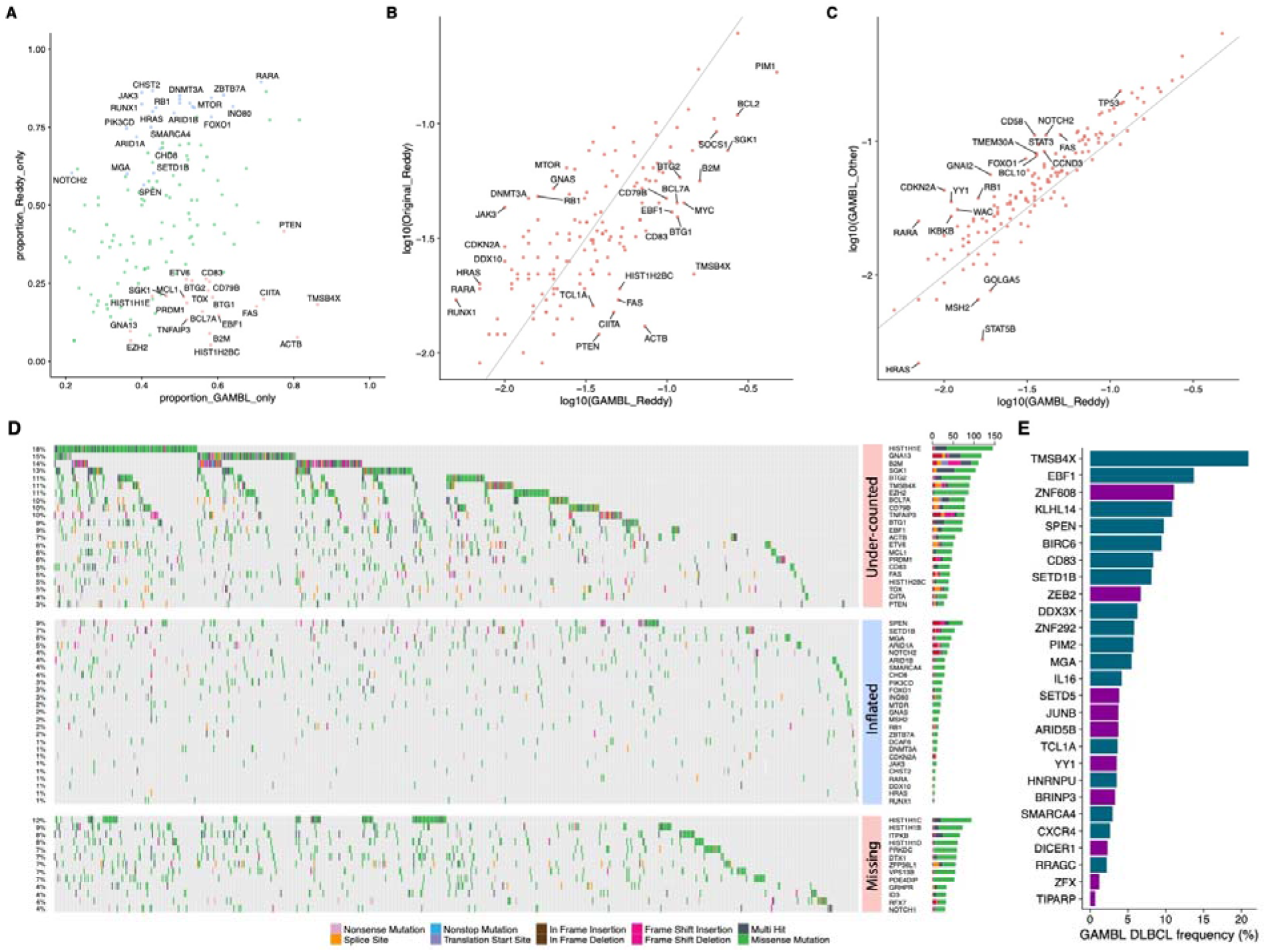
Mutation frequencies in candidate and established DLBCL genes. (A) The proportions of mutations in the GAMBL-only and Reddy-only sets are compared and genes with significantly different proportions are labeled. Blue points above the diagonal represent genes with a higher proportion of variants unique to the Reddy results. Assuming the GAMBL results are more accurate, these are the genes with the most severe inflation of mutations in the Reddy analysis. Salmon-coloured points below the diagonal represent the converse scenario, that is, genes with significantly higher mutation rates based on the GAMBL analysis. (B) The mutation frequency according to the GAMBL results and the Reddy results are compared. Similar to (A), the genes above the diagonal had inflated mutation incidence in the Reddy results whereas mutations were under-represented in the genes below the diagonal. These genes include a mix of genes with known hot spots (e.g. *EZH2*, *CD79B*, *B2M*) and genes affected by aSHM (e.g. *CD83*, *TMSB4X*, *CIITA*), indicating that these scenarios may have been inadequately handled by the Reddy analysis. (C) To assess whether the Reddy cohort provides a reasonable representation of mutations in DLBCL, the mutation incidence in the Reddy genes are contrasted between this cohort and 912 DLBCLs from a collection of other studies. Although there was a slight under-representation of mutations in all genes in the Reddy cases, four genes show the opposite trend, namely *GOLGA5*, *MSH2*, *HRAS* and *STAT5B*. (D) An oncoplot showing the mutation status of each patient based on the GAMBL results. The upper and lower sections respectively show the genes with mutations under-reported in the Reddy results and a selection of established DLBCL genes that were **absent** from the Reddy gene list. The middle section shows the mutation status of the genes that had significantly higher mutation rates according to Reddy. (E) The frequency of mutations (based on GAMBL) of 27 genes that were described as novel by Reddy. Dark blue indicates genes that we consider to be valid DLBCL drivers.

**Figure 3.**
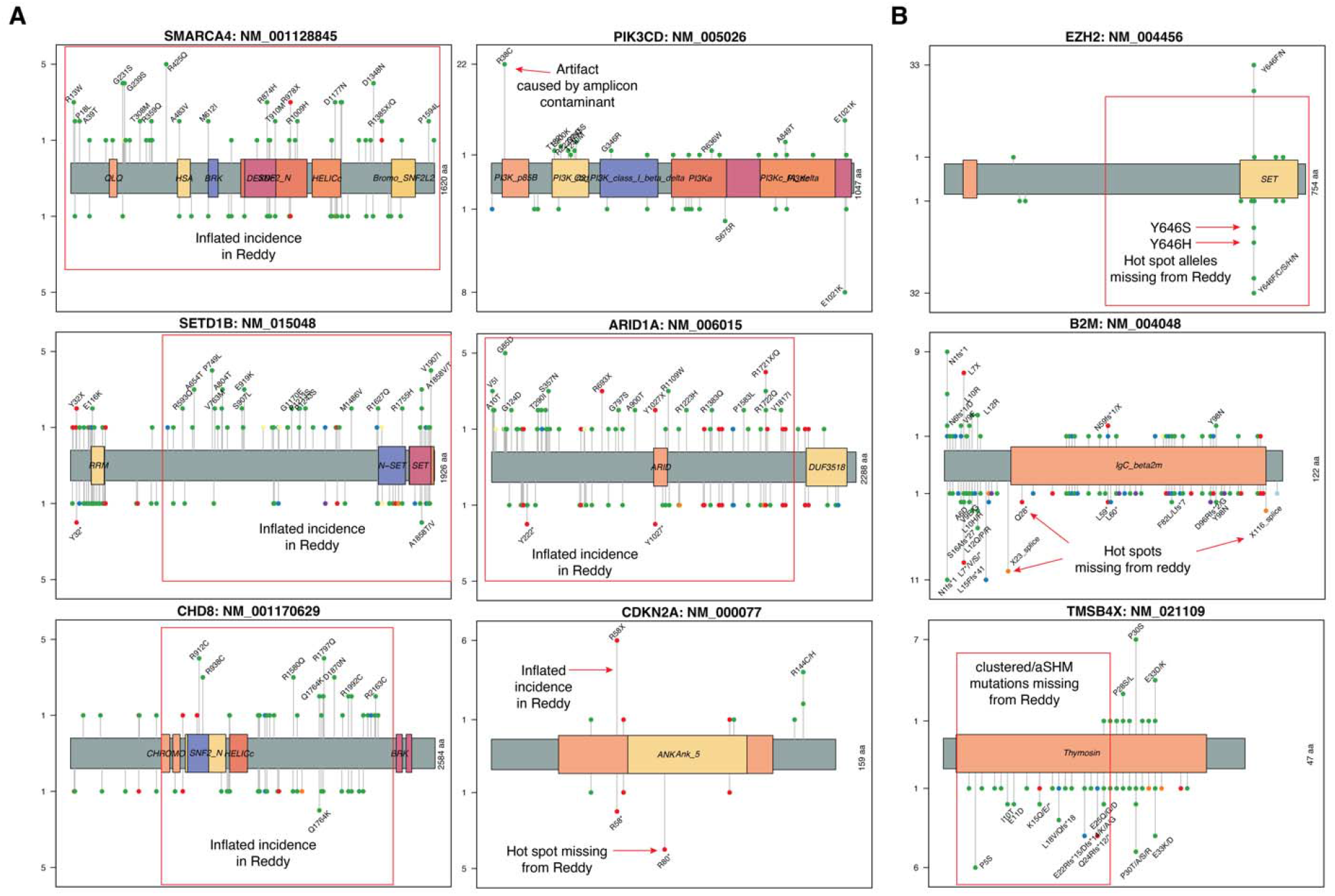
Contrasting mutation patterns between the Reddy and GAMBL results. Each lollipop plot shows the location of mutations and predicted effect on the protein for variants reported in the Reddy study (above) and the GAMBL results (below). The height represents the number of patient samples in which the mutation was detected. Regions of proteins with particularly discrepant mutation patterns are indicated with red arrows or boxes. (A) Six examples of genes with a substantial number of variants that were unique to the Reddy results (inflated incidence). (B) Three examples of genes with a substantial proportion of mutations in the GAMBL results that were not reported by Reddy.

The supplemental tables only reported the mutation status of 150 genes, thereby redacting all other mutations. In addition to the GAMBL-only list, which was restricted to the 150 Reddy genes, there were 2859 additional GAMBL variants detected in the Reddy exomes among additional DLBCL genes. This included genes that had higher mutation incidences than in any of the novel genes reported by Reddy. Thirteen of these had previously, or have since, been reported as recurrently mutated in DLBCL or other mature B-cell neoplasms (**Figure 2D, bottom**). For example, *HIST1H1C* and *HIST1H1B* were respectively mutated in 12% and 9% of patients. **Table 3** compares the mutation frequency of these and additional DLBCL genes in the Reddy samples and the remaining DLBCLs in GAMBL.

### Hot or not spots?

Genes with many GAMBL-only variants tended to be those affected by aberrant somatic hypermutation (aSHM) and other genes with highly recurrent mutation sites (hot spots)(**Figure 3**). *EZH2* has one major “canonical” mutation hot spot affecting codon 646.^10^ This mutation has four common alleles divided between two adjacent coordinates in the same codon (**Figure 4**). These alleles individually result in substitution of tyrosine with any of S, F, H or N. Only two of these alleles were reported in the Reddy results (F and N) with the other two conspicuously absent. Overall, the Reddy analysis failed to report mutations at this hot spot in 32 patients. Similarly, *CD79B* has a hot spot with multiple alleles at two adjacent coordinates and 15 examples of these mutations were missing from the Reddy results, again with only two alleles ever reported. The effect of this error on each gene was variable but, clearly, it caused them to underestimate the mutation prevalence in genes with common hot spots.

**Figure 4.**
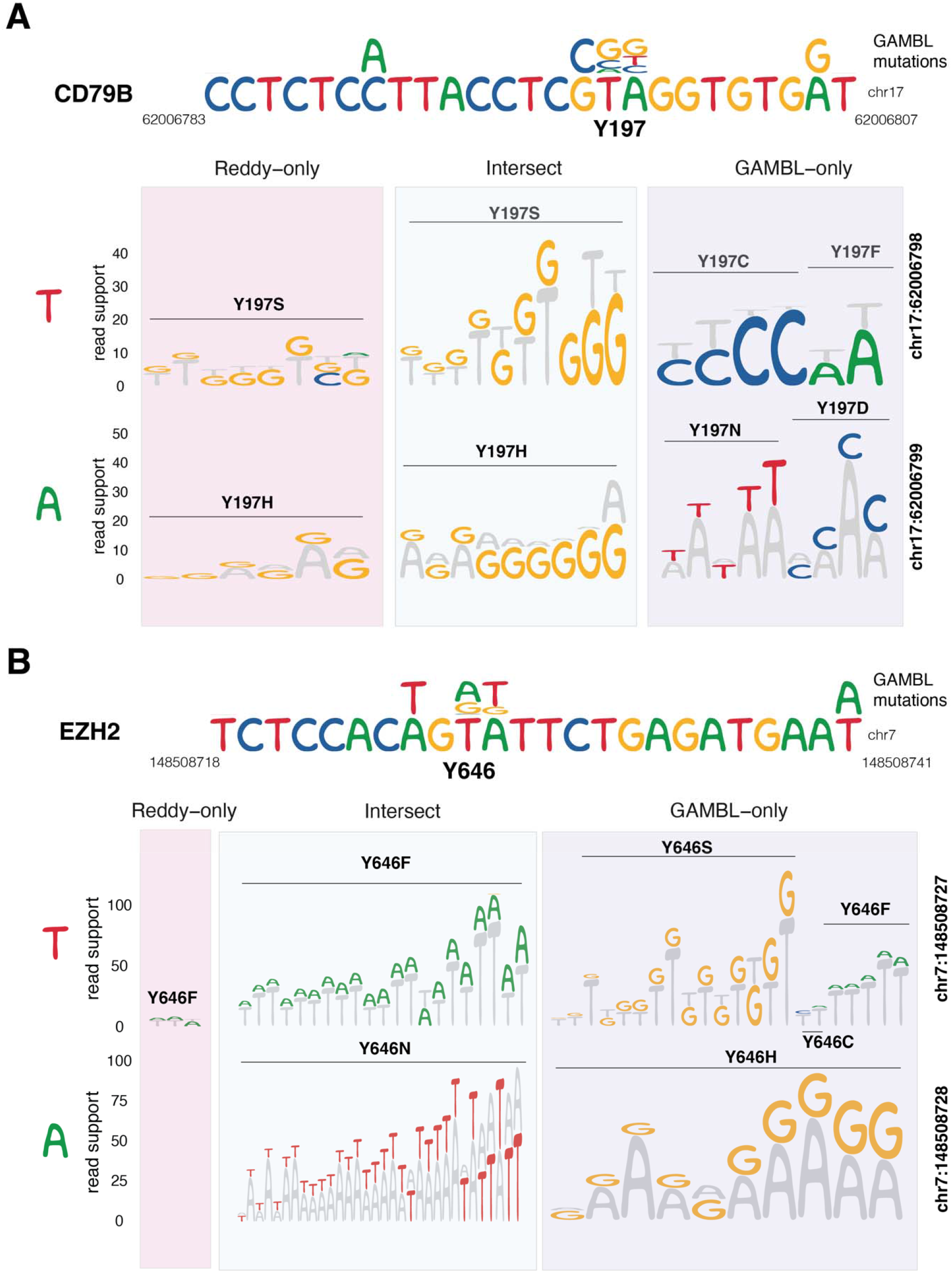
Systematic under-representation of mutation hot spots in Reddy results. Each panel shows the genomic context of a multi-allelic hotspot with the upper section showing all mutations found in the region within the GAMBL results. In both examples, the affected codon is GTA (TAC on the coding strand). The lower section shows the read support for the reference (gray) and non-reference bases in individual patients with mutations reported at each of the two coordinates either in the original Reddy analysis or GAMBL. (A) *CD79B* Y197 had two alleles reported in Reddy, one at each coordinate in the codon (T>G and A>G). There were additional examples of these mutations reported by Reddy that were not in our results (pink section). The GAMBL results include four additional alleles affecting this codon (purple section), totaling 15 patients. (B) Similarly, *EZH2* Y646 had two alleles reported by Reddy (T>A and A>T). There were very few mutations in this hot spot that were unique to the Reddy results and they each had low read support (pink section). Again, the GAMBL results included four additional mutant alleles that were not reported by Reddy, totaling 32 patients.

Using the GAMBL results, we naively searched for putative multi-allelic hot spots across the Reddy cohort, as defined by positions with at least two different substitution mutations observed in the cohort. This identifies 294 multiallelic hot spot sites across the 150 genes, collectively representing 1212 mutations in GAMBL. Overall, 644 mutations that were missing from the Reddy results could be explained by this phenomenon. In some cases, as in the examples above, the Reddy results reported only one allele at a hot spot whereas others were completely absent (e.g. ETV6 K11). Collectively, 53% of the mutations at multiallelic sites were missing from the Reddy results due to one (or more) errors in their analysis. The hot spots affected by the most substantial drop-out in the Reddy results are shown in **Table 2** with the entire set included in the accompanying supplement (**Supplemental Table S1**).

**Table 2.**
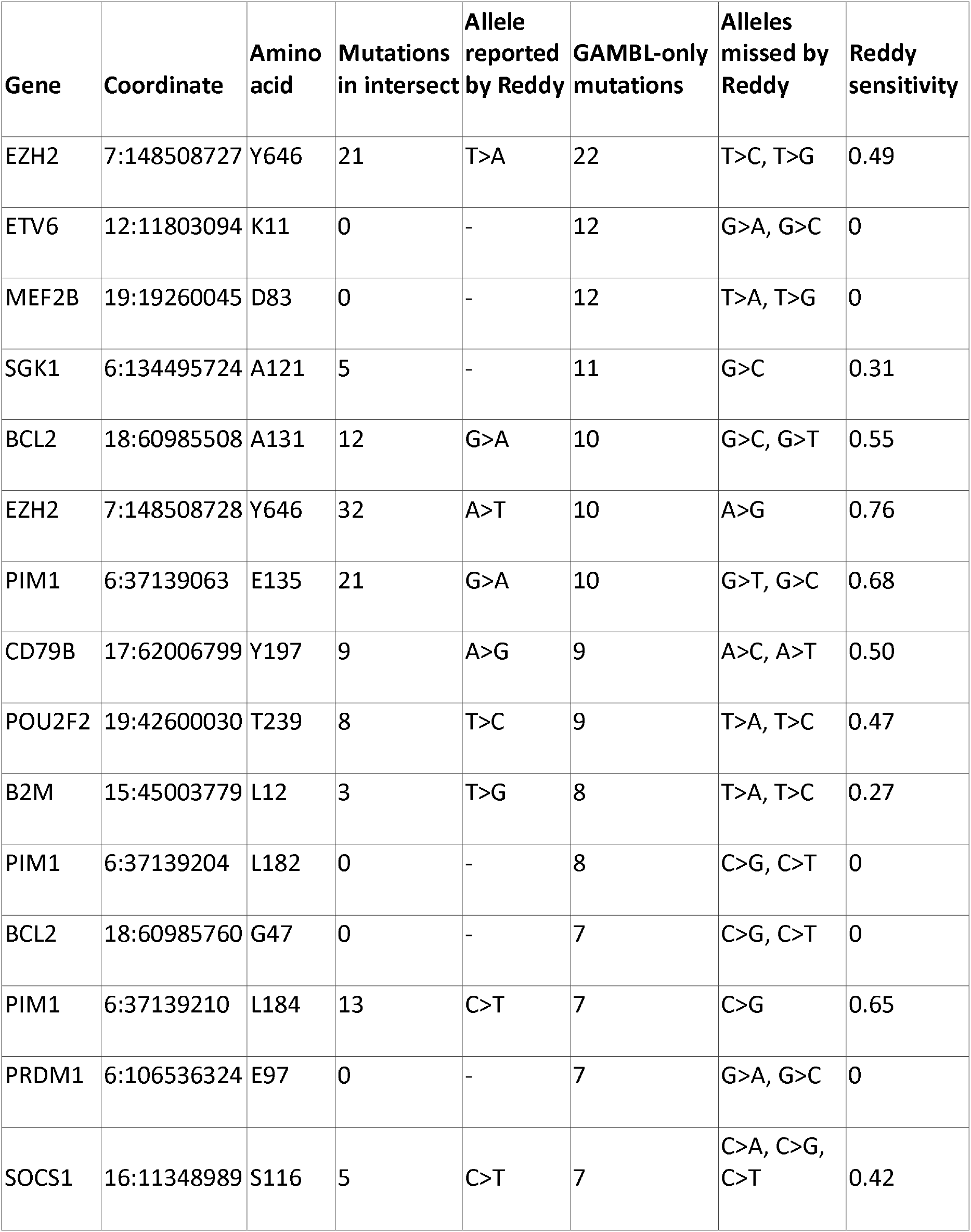
Multi-allelic hot spot mutations that were systematically dropped by the Reddy analysis.

### COO-recting discrepancies and filling in gaps

DLBCL can be divided into two molecular subgroups ABC and GCB based on gene expression information, with a smaller unclassifiable subset. Numerous studies have demonstrated an association between the mutation status of DLBCL genes and either of these molecular subgroups. Using COO labels (available for the 753 patients with RNA-seq data), the Reddy analysis identified 20 genes that were mutated more commonly in either ABC (8) or GCB (12) DLBCL. Using the GAMBL set of variants, we were able to confirm many of the associations presented by Reddy with three exceptions: *B2M*, *MGA* and *CDKN2A*.

The GAMBL results also identified additional COO-associated genes that were not among those reported by Reddy. In particular, *GRHPR* and *MPEG1* are enriched for mutations in ABC and *HVCN1*, *S1PR2*, *DDX3X*, more commonly mutated in GCB. **(Figure 5).** These discrepancies can be explained by a combination of factors. Each of *GRHPR*, *MPEG1* and *HVCN1* were not in the Reddy gene list. For the remaining two genes, we noted a higher mutation load in the GAMBL results, which may have provided increased power to detect these differences. *S1PR2* and *DDX3X* respectively had 3 and 13 additional patients with mutations in the GAMBL analysis. Notably, most of these genes are important features in the genetic subgrouping systems such as LymphGen and clearly belong on the DLBCL gene list.^11^

**Figure 5.**
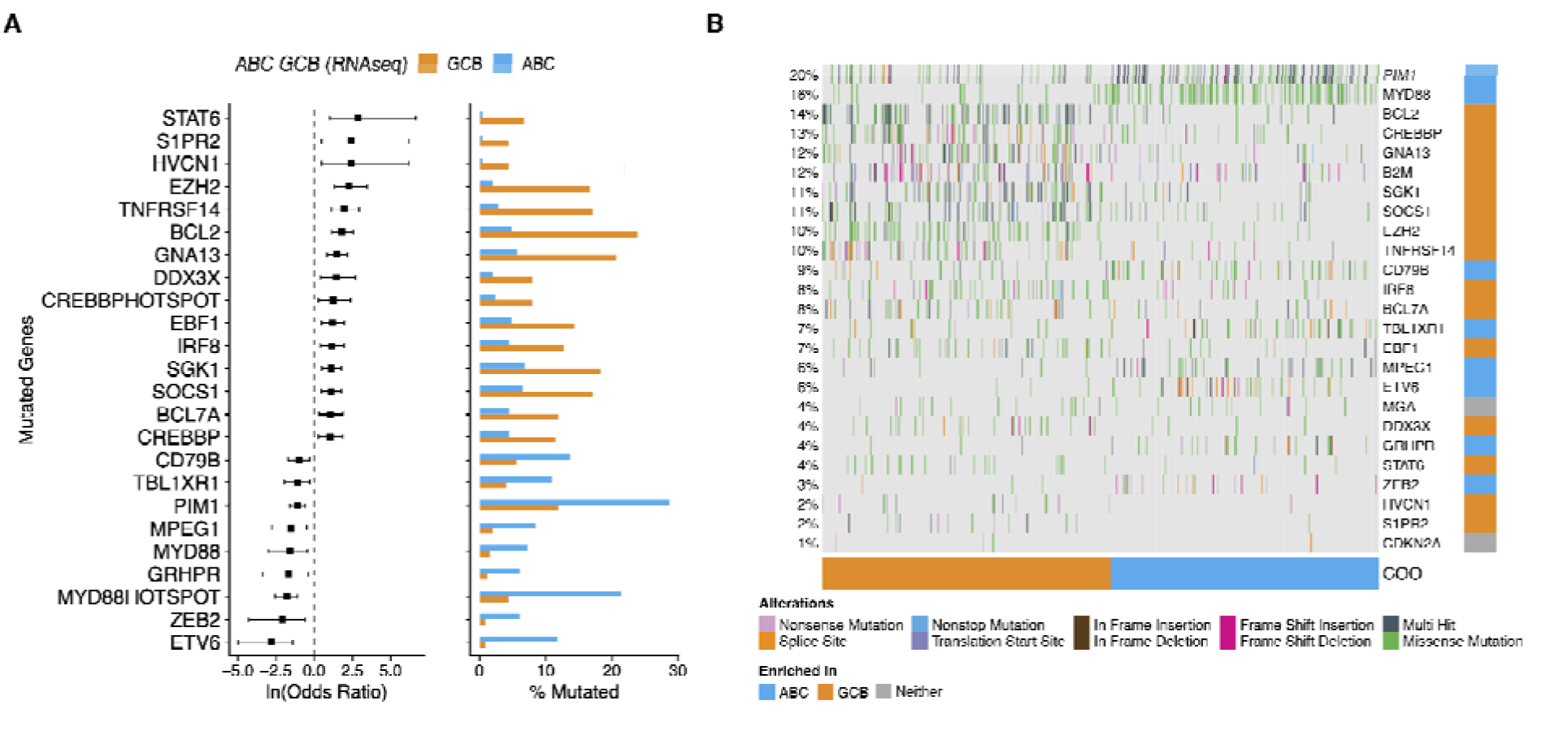
Genes with mutations associated with ABC or GCB DLBCL. (A) The GAMBL results identified more genes with mutations associated with one of the two molecular subgroups. The genes that were individually significant (Fisher’s Exact test, P < 0.05) are shown alongside their mutation incidence in each subgroup based on GAMBL. (B) The mutations identified in these genes across all ABC and GCB DLBCLs are shown along with the two genes for which the association reported by Reddy could not be reproduced (*MGA* and *CDKN2A*).

### Most survival correlates did not survive

Reddy demonstrated associations between mutation status and overall survival for 5 genes. In a subgroup analysis within the ABC and GCB cases, they identified another 10 genes associated with outcomes. Using the entire cohort, they reported negative prognostic associations with *MYC*, *CD79B* and *ZFAT* and positive prognostic associations with *NF1* and *SGK1* mutations. Using the GAMBL results, we could only reproduce the negative prognostic association with *MYC* mutations. The loss of the additional associations can be explained by the large number of discrepancies in identified mutations among these genes. For example, the GAMBL analysis only identified 11 patients with NF1 mutations, in contrast to the 33 patients that were mutated according to Reddy. Repeating their analysis with the GAMBL variant calls also allowed us to detect significant negative prognostic associations with mutations in *TP53*, *KMT2D*, *PIM1*, and *PDE4DIP* (**Figure 6A**). *KMT2D* mutation status was also associated with reduced overall survival times in the GCB subset and *TP53* status remained significant in both subsets (**Figure 6B**). Although none of these associations were reported by Reddy, we found that the association between *PIM1* and poor outcome was also highly significant with their mutation calls. In contrast, stratifying patients according to *TP53* or *KMT2D* mutation status using the Reddy results showed no difference in outcomes (**Figure 6C**).

**Figure 6.**
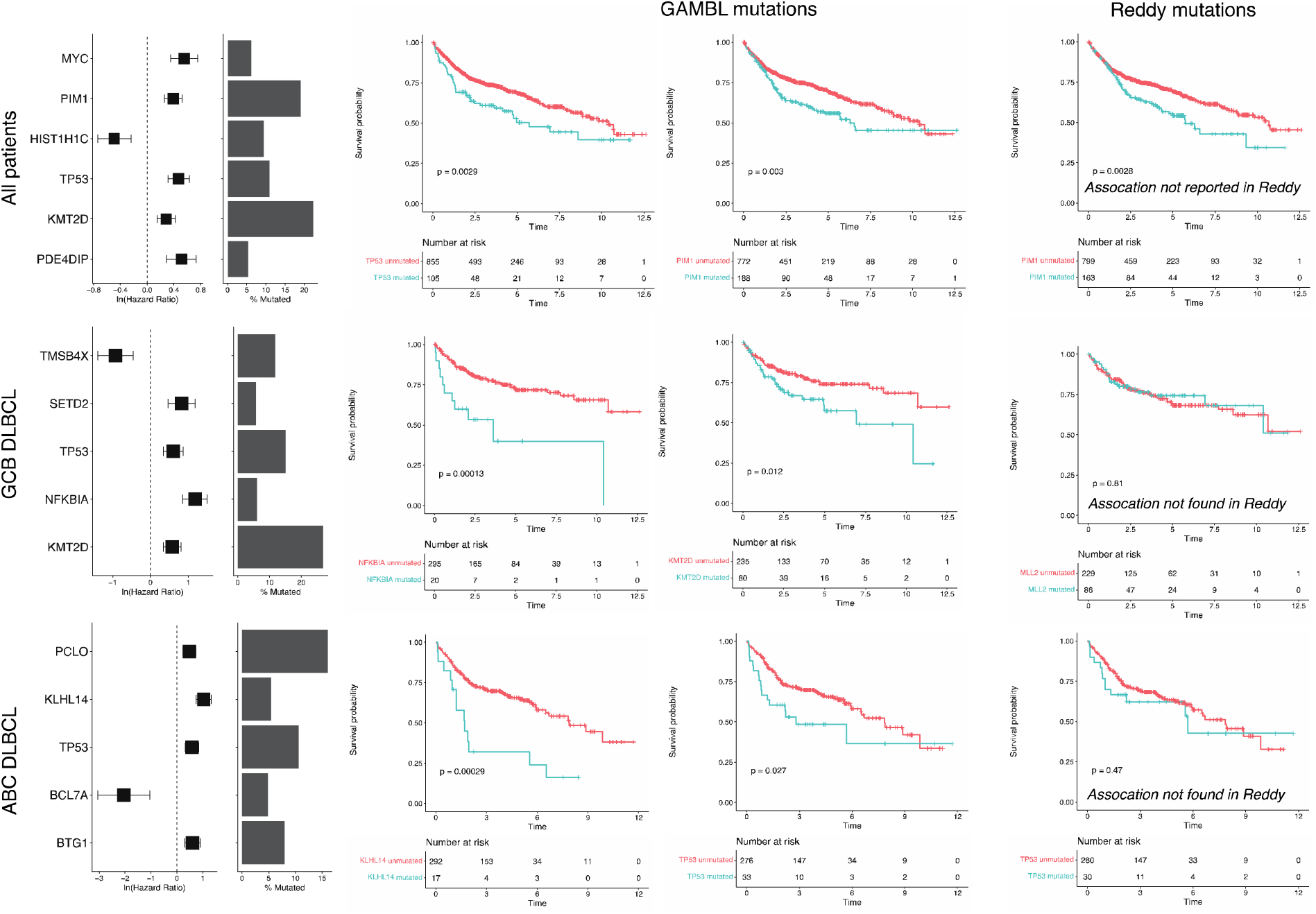
Associations between mutation status and patient survival. (A) The genes with significant associations with overall survival (OS) time using the entire cohort (top) or in either COO subgroup (middle and bottom) are shown as forest plots. With the hazard ratio and 95% confidence intervals shown (on a log scale). The proportion of cases mutated is also included as a bar chart. Most of the associations we detected were not found by Reddy (genes indicated in red) (B) Selected associations were represented as Kaplan-Meier curves stratifying on the mutation status according to the GAMBL results. (C) For comparison, we generated Kaplan-Meier curves for three genes using the Reddy mutations. Interestingly, PIM1 shows a strong negative prognostic association with both sets of mutations but this association was not reported by Reddy. The associations for *KMT2D* and *TP53* are not reproducible using the Reddy mutations.

## Discussion

### Cause

The magnitude of discrepancies between the GAMBL mutations and those presented in the original study is astonishing. To understand the effect of these discrepancies we must consider them individually. The Reddy-only variants could mostly be assigned to one of two categories. The bulk of those variants had minimal supporting evidence in the primary data, as determined by visual review and variant allele fraction. The cause of these variants can be easily explained by carefully revisiting the methods described by the authors. They vaguely describe a joint variant calling technique in which they use the existence of a mutation in one patient as prior evidence for that mutation in all other patients. **Figure 7** shows how this approach leads to the artificial creation of apparent mutation hot spots, often with low read support. As a result of this strategy, the Reddy-only variants had generally low read support (mean VAF = 0.256) compared to the Intersect (mean VAF = 0.420). Although this form of variant calling is suited for germline applications, it is inappropriate to apply this to cancer data because hot spots are the exception rather than the rule. The remaining Reddy-only variants tended to have strong support in the sequence data along with gnomAD allele frequencies that were highly suggestive of them originating in the germline rather than representing somatic mutations (**Figure 1D-E**). A small number of Reddy-only variants did not fit in either category, instead appearing to represent highly recurrent artifacts caused by contamination of the exomes with amplicon. Collectively, these variants caused an artificial inflation of the mutation incidence in many genes including several of the novel DLBCL genes presented in this study.

**Figure 7.**
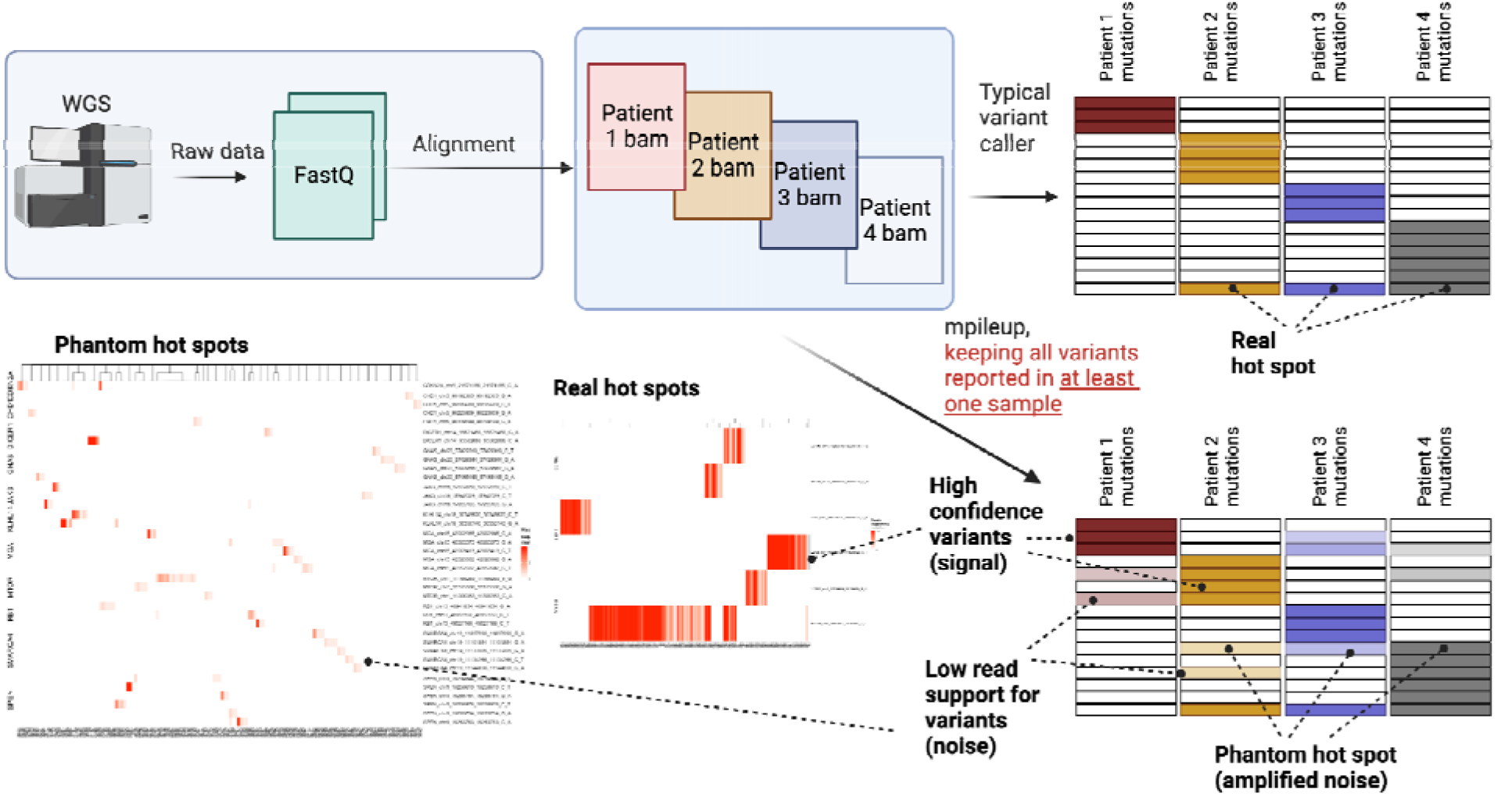
Joint variant calling promotes phantom hot spots. Reddy describes two separate approaches for detecting somatic mutations. They used MuTect to identify variants in the cases having a matched normal exome. They also describe using “joint variant calling” on all 1001 samples using samtools mpileup. They also state “each variant was required to have an instance of genotype quality greater than 30 and read depth greater than 5” This implies that the authors considered a variant to be valid in any patient (regardless of the read support) if that same allele was detected in at least one other patient. This leads to error propagation, allowing the same variant to be “detected” in more than one patient due to sequencing errors or artifacts that occur at that position by coincidence.

The failure to report more than 3000 additional high-quality variants among the Reddy genes was equally concerning. Although the absence of these cannot be explained by a single phenomenon, some patterns have emerged that inform on possible underlying causes. Ironically, although the strategy used by Reddy appears to have created numerous phantom hot spots, the same approach seems to have caused the suppression of hundreds of *bona fide* hot spots across 54 genes (**Table 2; Supplemental Table S4**). Specifically, their use of *samtools mpileup* to identify variants in the entire cohort in unison would result in a single VCF with multiallelic variants at each of those positions. As explained in the Annovar documentation, multiallelic variants must be split prior to annotation (using *bcftools norm*). The most likely explanation for the missing hotspots was that this step was not performed. This would effectively reduce every such variant to a single allele, thereby blinding them to all other alleles in their results. As mentioned previously, hot spots are not particularly common in cancer sequencing studies. Accordingly, most of the GAMBL-only variants remain unexplained.

### Consequence

The Reddy-only mutations were not uniformly distributed across genes. **Figure 2E** highlights the revised mutation frequency of the 27 Reddy genes that were presented as novel DLBCL genes. In our results, the mutation frequencies of half of these are below 5%. Our team has established a DLBCL gene list that has been curated using the available data from all large exome- and whole genome sequencing studies. Some additional genes with provisional status in our curated DLBCL list were similarly impacted by the artificial inflation identified herein such as *CHST2*, *JAK3*, *RUNX1*, *PIK3CD*, *DNMT3A*, *RARA*, *ZBTB7A*, *INO80*, and *HRAS* (**Figure 2A, D**). Considering the more accurate, albeit conservative, GAMBL results, the mutation incidence of those genes ranges from 1% and 4% in the Reddy cohort, again drawing their relevance in DLBCL into question. Due to the convoluted approach employed by Reddy, involving in-house scripts that were also used in their earlier studies, we are unable to evaluate whether a similar analysis would prioritize these same 150 genes. It seems unlikely, though, given the existence of genes with mutation rates as high as 12% that were not identified by their analysis. In summary, although we agree there may be approximately 150 DLBCL genes, based on our analysis, numerous important genes were missing from the Reddy gene list and equally as many on that list that likely have no relevance to DLBCL.

While mutation prevalence in some genes was inflated by the Reddy analysis, their results also under-reported the prevalence of hundreds of mutation hot spots. In some cases, Reddy reported only the most common allele but in others, the hot spot was completely redacted (**Table 3**; **Supplemental Table S4**). The existence of some of these mutations was already firmly established at the time of the Reddy study, for example those shown in **Figure 4**. *CD79B* ITAM mutations were one of the original genetic features associated with the dependence of ABC DLBCL on B-cell receptor signaling, a feature that has been exploited through BTK inhibitors such as Ibrutinib.^12,13^ Whereas the Reddy analysis reported *CD79B* mutations in 4.7% of their cohort, we found them in 10% of the Reddy cases and 16.9% of the non-Reddy DLBCLs. Similarly, in Follicular lymphoma (FL) and DLBCL, EZH2 Y641 mutations promote its enzymatic activity, leading to an accumulation of H3K27me3 at polycomb repressive complex 2 (PRC2) targets. EZH2 inhibitors have been shown to provide therapeutic benefit in FL patients and continue to be explored in DLBCL.^5,14,15^ Reddy reported an *EZH2* mutation frequency of 6.1%, whereas 9.4% of the Reddy cases and 9.6% of the non-Reddy DLBCLs are mutated according to GAMBL.

**Table 3.** DLBCL genes not reported in Reddy. The genes with mutation frequency of at least 4% in GAMBL are shown along with the mutation frequency in GAMBL DLBCLs (non-Reddy) and the GAMBL Reddy DLBCLs. Whether each gene was considered significantly mutated in other DLBCL studies is indicated with citations. An asterisk indicates a significantly lower frequency in Reddy compared to the other DLBCLs in GAMBL.

| Gene | mutation frequency (other DLBCL) (%) | mutation frequency (Reddy cases)(%) |
| --- | --- | --- |
| DTX1<sup>2,311/21/2023</sup> | 17.3 | 6.9* |
| MPEG1<sup>3</sup> | 15.6 | 10.5* |
| HLA-B<sup>2,3</sup> | 15.5 | 1.4* |
| HIST1H1C<sup>10</sup> | 13.5 | 11.8 |
| ITPKB<sup>3</sup> | 12.1 | 10.5 |
| OSBPL10<sup>3</sup> | 12.0 | 2.5* |
| HLA-A<sup>2,3</sup> | 11.6 | 0.2* |
| ZFP36L1<sup>2,3</sup> | 11.3 | 6.7* |
| HIST1H1B<sup>2</sup> | 10.5 | 8.4 |
| KLF2<sup>3</sup> | 8.8 | 2.5* |
| HIST1H1D<sup>2</sup> | 7.9 | 6.5 |
| HIST1H2AC<sup>2</sup> | 7.2 | 3.9* |
| PRKDC<sup>3</sup> | 7.0 | 7.1 |
| HIST1H2AM<sup>2</sup> | 6.4 | 4.9 |
| HIST1H2BC<sup>2</sup> | 5.9 | 5.1 |
| HIST1H2BK<sup>2</sup> | 5.7 | 4.6 |
| NOTCH1<sup>3</sup> | 5.7 | 3.9 |
| ZC3H12A<sup>2</sup> | 5.7 | 3.5* |
| GRHPR<sup>2,3</sup> | 5.6 | 4.8 |
| NOL9<sup>3</sup> | 5.2 | 1.4* |
| BCL11A<sup>2</sup> | 4.6 | 1.6* |
| ID3<sup>3</sup> | 4.3 | 4.3 |
| FOXC1<sup>3</sup> | 4.1 | 1.5* |

Beyond the relevance of individual mutations in precision medicine, we note that many of the mutations with discrepancies between the Reddy and GAMBL results contribute to DLBCL genetic subgroup assignments, such as LymphGen.^11^ The Reddy analysis was particularly inadequate at detecting mutations in genes affected by aSHM (**Figure 2A**). This may relate to the issue that caused multi-allelic variants to be dropped and/or could be explained by another problem in their analysis. Regardless of the underlying cause, the loss of any mutations associated with genetic subgroups would clearly reduce the accuracy of the resulting classification. The importance of accurate and complete mutation detection in all such genes will become paramount as genetic subgroups begin to be incorporated into clinical trials.^6,15^

Beyond delineating the individual genes and mutation patterns relevant to DLBCL, the Reddy study presented a host of associations between mutation status and clinical features including molecular subgroup (COO) and patient survival (“Genomic Risk”). In contrast to their results, we did not find an association between *MGA*, *B2M* or *CDKN2A* mutation status and COO. Some of these discrepancies might be attributable to the use of copy number information in the Reddy study, which was not used in this re-analysis. Indeed, 86 patients were assigned as *CDKN2A*-mutated by Reddy and only 30 of these represent patients assigned a SSM by Reddy. Unfortunately, the majority of *CDKN2A* mutations were not reproduced in the GAMBL analysis and are likely enriched for artifacts (**Figure 3**). Surprisingly, the most abundant *CDKN2A* mutation in the GAMBL results (R80X) – also the most common mutation in COSMIC – was not reported by Reddy. Notably, the association between deletions of the *CDKN2A* locus and ABC DLBCL was established long before this study^16^ but an association with SSM (i.e. the aspect we could not reproduce), would have been a novel contribution to the literature. Instead, our results indicate that additional genes, not reported in their study, are significantly enriched for mutations in ABC or GCB DLBCL. These differences can be attributed to the large number of discordant mutations between the two analyses.

Finally, our analysis showed that most of the associations between mutation and patient outcomes presented in the original study are not reproducible. Such shifts in results were not particularly surprising considering the magnitude of the differences between the GAMBL and Reddy variants. With the marked differences observable with univariate analyses, it appears unlikely that the Genomic Risk Model would prove to be robust. Given the newly identified associations with commonly mutated genes such as *TP53*, *PIM1* and *KMT2D*, a risk model using the GAMBL results should clearly outperform any model derived from the original variants.

## Experimental Procedures

### Statistical analysis and data visualization

Custom analyses and data visualizations were performed in R, primarily using the GAMBLR package (https://github.com/morinlab/GAMBLR). Specifically, COO associations with mutation status were performed using “prettyForestPlot” function. All oncoplots were generated with “prettyOncoplot” function (**Figure 2**, **Figure 5**). Lollipop plots (**Figure 3**) were generated using MAFtools (https://bioconductor.org/packages/release/bioc/html/maftools.html). Kaplan-Meier plots were generated using survminer (https://cran.r-project.org/web/packages/survminer/index.html). Mutations were visualized as sequence logos (**Figure 4**) using the ggseqlogo package (https://omarwagih.github.io/ggseqlogo/).

### Variant calls

For comparison purposes, we have also assembled a cohort of 912 of DLBCL comprising tumors that underwent whole-genome or whole-exome sequencing and comprehensively characterized previously by several independent groups. Specifically, the comparison cohort in GAMBL consisted of 703 DLBCLs analyzed by whole exome sequencing in the studies by Schmitz et al and Chapuy et al, as well as 209 DLBCLs with whole-genome sequencing from the studies by Morin et al^17^, Arthur et al^18^, Hilton et al^19^, and the MMML-seq project. For compatibility with the GAMBL pipeline, numeric sample ID from each study was modified following the convention of prefixing each with study’s first author and adding suffix T to indicate tumor. For example, sample 1008 from the Reddy study became Reddy_1008T.

Tumor exomes were analyzed for somatic mutations using a combination of four modern variant callers, an approach that we have previously shown to perform favorably on data from formalin-fixed paraffin embedded (FFPE) tissue.^20,21^ The variants reported by Reddy were converted from the wide tabular format into a long MAF-like tabular format for subsequent analyses. The read support for each variant was added using a custom Python script that relies on pysam (https://github.com/LCR-BCCRC/lcr-scripts/tree/master/augment_ssm/1.0). To identify the Reddy-only, GAMBL-only and intersect lists, we compared the SNVs reported by the authors to the variants obtained by applying our pipeline to the 999 exome samples that were available. Matching of variants between this table and the GAMBL MAF was performed using chromosome, position, reference allele, alternate allele and sample ID. Variants were annotated using the Variant Effect Predictor and vcf2maf and all variants with gnomAD AF > 0.0001 were removed.

### Variant quality assessment

Variants in all lists were subjected to visual review using a set of images generated by IGV. For each variant, the raw data (bam) was loaded into IGV and a snapshot was saved with sufficient dimensions to ensure the reads in the surrounding region were visible, generally representing the window of 150 bases. Visualizations were generated with reads in paired view as well as separate and with/without soft-clipped portions of reads shown. These images were available to curators through a custom web interface implemented in R using Shiny. For each variant, the interface recorded the rating, user ID, a set of optional tags, and optional freeform comments. The rating scale and tags were derived from a standard operating procedure with modifications to account for the unavailability of matched normal samples from the Reddy study. Most of the variants were rated by four curators. These individuals were, in descending order of the number of variants rated: HL, NL, RM, and CB.

### Hot spot analysis

Multi-allelic hot spots were identified from the GAMBL results by grouping all variants by genomic coordinate and identifying coordinates with at least two distinct non-reference alleles. As described in the Reddy study, we used the samtools *mpileup* command to summarize the support for mutations in each of these regions (using all 999 bam files as input). These outputs were used to determine the support for each non-reference allele identified by GAMBL including those missing from the Reddy results.

## Acknowledgements

This study relies on the raw data from Reddy et al, which was obtained from EGA (EGAD00001003600) with approval from the senior author of that study. The authors also acknowledge the ICGC MALY-DE project (https://dcc.icgc.org) for providing access to their data. Aligned reads for those genomes were obtained through a DACO-approved project (R.D.M.) using a virtual instance on the Cancer Genome Collaboratory. The results published here are in whole or in part based upon data generated by the Cancer Genome Characterization Initiative (phs000235), Non-Hodgkin lymphoma and whole exome sequencing of diffuse large B-cell lymphoma (phs000450), developed by the National Cancer Institute. Some of the data used for these analyses are available from dbGAP accessions phs000235.v14.p2, phs000527.v3.p1, and phs000450.v1.p1. All data were used according to the data use agreements.

## References

1. Reddy, A. et al. Genetic and Functional Drivers of Diffuse Large B Cell Lymphoma. Cell 171, 481–494.e15 (2017).

2. Chapuy, B. et al. Molecular subtypes of diffuse large B cell lymphoma are associated with distinct pathogenic mechanisms and outcomes. Nat. Med. 24, 679–690 (2018).

3. Schmitz, R. et al. Genetics and Pathogenesis of Diffuse Large B-Cell Lymphoma. N. Engl. J. Med. 378, 1396–1407 (2018).

4. Petereit, J. Pipeline Automation via Snakemake. Methods Mol. Biol. Clifton NJ 2443, 181– 196 (2022).

5. Morin, R. D., Arthur, S. & Assouline, S. Treating lymphoma is now a bit EZ-er. Blood Adv. 5 8, 2256–2263 (2021).

6. Mondello, P. & Ansell, S. PHOENIX rises: Genomic-based therapies for diffuse large B cell lymphoma. Cancer Cell (2021) doi:10.1016/j.ccell.2021.10.007.

7. Barnell, E. K. et al. Standard operating procedure for somatic variant refinement of sequencing data with paired tumor and normal samples. Genet. Med. 21, 972–981 (2019).

8. Robinson, J. T. et al. Integrative genomics viewer. Nat Biotechnol 29, 24–26 (2011).

9. Zhang, J. et al. Genetic heterogeneity of diffuse large B-cell lymphoma. (2013).

10. Morin, R. D. et al. Somatic mutations altering EZH2 (Tyr641) in follicular and diffuse large B-cell lymphomas of germinal-center origin. Nat. Genet. 42, 181–185 (2010).

11. Wright, G. W. et al. A Probabilistic Classification Tool for Genetic Subtypes of Diffuse Large B Cell Lymphoma with Therapeutic Implications. Cancer Cell 37, 551–568.e14 (2020).

12. Davis, R. E. et al. Chronic active B-cell-receptor signalling in diffuse large B-cell lymphoma. Nature 463, 88–92 (2010).

13. Brown, J. R. PCI-32765, the First BTK (Bruton’s Tyrosine Kinase) Inhibitor in Clinical Trials. Curr. Hematol. Malig. Rep. 8, 1 (2013).

14. Mt, M. et al. EZH2 inhibition as a therapeutic strategy for lymphoma with EZH2-activating mutations. Nature 492, (2012).

15. Zhang, M. et al. Genetic subtype-guided immunochemotherapy in diffuse large B cell lymphoma: The randomized GUIDANCE-01 trial. Cancer Cell (2023) doi:10.1016/j.ccell.2023.09.004.

16. Lenz, G. et al. Molecular subtypes of diffuse large B-cell lymphoma arise by distinct genetic pathways. Proc. Natl. Acad. Sci. 105, 13520–13525 (2008).

17. Morin, R. D. et al. Mutational and structural analysis of diffuse large B-cell lymphoma using whole-genome sequencing. Blood 122, 1256–1265 (2013).

18. Arthur, S. E. et al. Genome-wide discovery of somatic regulatory variants in diffuse large B-cell lymphoma. Nat. Commun. 9, 4001 (2018).

19. Hilton, L. K. et al. The double-hit signature identifies double-hit diffuse large B-cell lymphoma with genetic events cryptic to FISH. Blood 134, 1528–1532 (2019).

20. Dreval, K. et al. Genetic subdivisions of follicular lymphoma defined by distinct coding and noncoding mutation patterns. Blood 142, 561–573 (2023).

21. Thomas, N. et al. Genetic Subgroups Inform on Pathobiology in Adult and Pediatric Burkitt Lymphoma. 2021.12.05.21267216 Preprint at 10.1101/2021.12.05.21267216 (2021).

